# Metabolic Stability of the Demyelination PET Tracer [^18^F]3F4AP and Identification of its Metabolites

**DOI:** 10.1101/2022.09.27.509607

**Authors:** Yang Sun, Karla Ramos-Torres, Pedro Brugarolas

## Abstract

[^18^F]3-fluoro-4-aminopyridine ([^18^F]3F4AP) is a PET tracer for imaging demyelination based on the multiple sclerosis drug 4-aminopyridine (4AP, dalfampridine). This radiotracer was found to be stable in rodents and nonhuman primates imaged under isoflurane anesthesia. However, recent findings indicate that its stability is greatly decreased in awake humans and mice. Since both 4AP and isoflurane are metabolized primarily by cytochrome P450 enzymes, particularly CYP2E1, we postulated that this enzyme may be responsible for the metabolism of 3F4AP. Here, we investigated the metabolism of [^18^F]3F4AP by CYP2E1 and identified its metabolites. We also investigated whether deuteration, a common approach to increase the stability of drugs, could improve its stability. Our results demonstrate that CYP2E1 readily metabolizes 3F4AP and its deuterated analogues and that the primary metabolites are 5-hydroxy-3-fluoro-4-aminopyridine and 3-fluoro-4-aminopyridine N-oxide. Although deuteration did not decrease the rate of the CYP2E1 mediated oxidation, our findings explain the diminished *in vivo* stability of 3F4AP compared to 4AP and further our understanding of when deuteration may improve the metabolic stability of drugs and PET ligands.

**Significance Statement:** Understanding the metabolic stability of PET tracers is paramount to its application in humans as metabolism, which varies from person to person, can affect the target-to-background signal. This study identified the predominant enzyme that metabolizes the demyelination PET tracer [^18^F]3F4AP and its metabolites. These findings may allow assessment of whether the radiometabolites can get into the brain and potentially lead to tracers with enhanced stability. Furthermore, this study furthers our understanding of when deuteration can improve metabolic stability.

## Introduction

[^18^F]3-fluoro-4-aminopyridine ([^18^F]3F4AP, [^18^F]**2**) is a PET radiotracer for imaging demyelination based on the multiple sclerosis drug 4-aminopyridine (4AP, dalfampridine, **1**) (Fig. 1) (Brugarolas et al. 2018). Previous PET imaging studies with [^18^F]3F4AP in rodents and nonhuman primates have shown that this tracer is highly sensitive to demyelinated lesions in the brain (Brugarolas et al. 2018; Guehl et al. 2021), which has led to the recent translation to human studies (ClinicalTrials.gov identifier: NCT04699747). In monkeys scanned under isoflurane anesthesia, the tracer was found to be very stable (>90% parent fraction remaining 2h post-injection) with good reproducibility (Guehl et al. 2021). However, recent findings in humans scanned without anesthesia show that less than 50% parent remains in circulation 1h post intravenous administration (Brugarolas et al. 2022) and a study in mice that received [^18^F]3F4AP awake or under isoflurane showed that non-anesthetized mice readily metabolized the tracer whereas anesthetized animals did not (72±4% vs. 20±5% parent remaining in plasma after 35 min), (Ramos-Torres et al. 2022). The observed differences in stability between awake and anesthetized subjects strongly suggest that the high stability in anesthetized animals is mediated by isoflurane.

**Figure 1.**
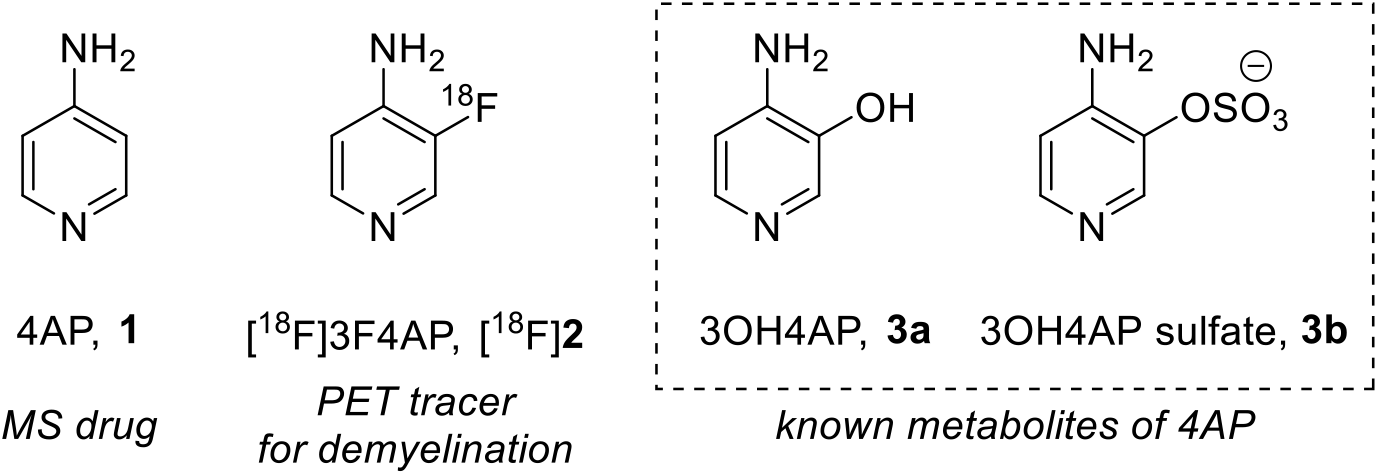
Structures of [_18_F]3F4AP, 4AP and known metabolites of 4AP.

A likely candidate responsible for the metabolism of [^18^F]3F4AP is the cytochrome P450 enzyme CYP2E1. CYP2E1 is the most abundant P450 enzyme in the human liver (Drozdzik et al. 2018) and is a key player in the breakdown of many small molecule drugs, including 4AP and isoflurane (Chen et al. 2019; Kharasch et al. 1999). The metabolism of 4AP has been investigated in rats, dogs, and humans (Caggiano et al. 2013; Caggiano and Blight 2013). These studies showed slow metabolism of 4AP, with unmetabolized 4AP amounting to at least 85% of the administered dose in humans 24 h after either oral or intravenous administration (Uges et al. 1982; Evenhuism et al. 1981). These studies also identified the primary metabolites as 3-hydroxy-4-aminopyridine (3OH4AP, **3a**) and the conjugated 3-hydroxy-4AP sulfate (**3b**) (Fig. 1) (Caggiano et al. 2013). The apparent ability of isoflurane to competitively inhibit the metabolism of [^18^F]3F4AP in rodents and primates further supports the role of CYP2E1 in 3F4AP metabolism.

Deuteration, a proven strategy in medicinal chemistry for increasing metabolic stability of bioactive compounds, could pose as a potential approach to inhibit the metabolism of 3F4AP. The difference in atomic mass of deuterium makes carbon-deuterium (C-D) bonds harder to break than corresponding carbon-hydrogen (C-H) bonds, thus decreasing the reaction kinetic rate for deuterated compounds compared to their nondeuterated counterparts. (Pirali et al. 2019; Gupta 2017; Wiberg 1955). This outcome can be quantitively expressed as the deuterium kinetic isotope effect (KIEs, i.e., k_H_/k_D_). Most reported examples of reactions catalyzed by cytochrome P450s show KIE values greater than 5 (Nelson and Trager 2003; Guengerich 2017). Although most of the examples of KIE pertain to the deuteration of aliphatic carbons, there is at least one reported example in which the deuteration of an aromatic compound (chlorobenzenes) resulted in a reduced reaction rate by P450 (Korzekwa et al. 1989).

This approach has also been used in PET to reduce the presence of radiometabolites, which can result in off-target signals (Ghosh et al. 2020; Lai et al. 2021; Raaphorst et al. 2018; Terry et al. 2010; Smith et al. 2011; Schou et al. 2004; Kuchar and Mamat 2015; Fowler et al. 1995; Xiao et al. 2021). In several cases, deuteration has successfully reduced defluorination and the accumulation of PET signals in bone (Lai et al. 2021; Raaphorst et al. 2018; Terry et al. 2010; Schou et al. 2004) (Fig. 2, **4-7**). Additionally, this deuteration strategy has contributed to the elimination of high background signal introduced by the oxidized radiometabolites of nondeuterated PET tracer **8**. (Smith et al. 2011) (Fig. 2, **8**). Given these findings, we hypothesized that deuterated 3F4AP may be more stable than nondeuterated 3F4AP.

**Figure 2.**
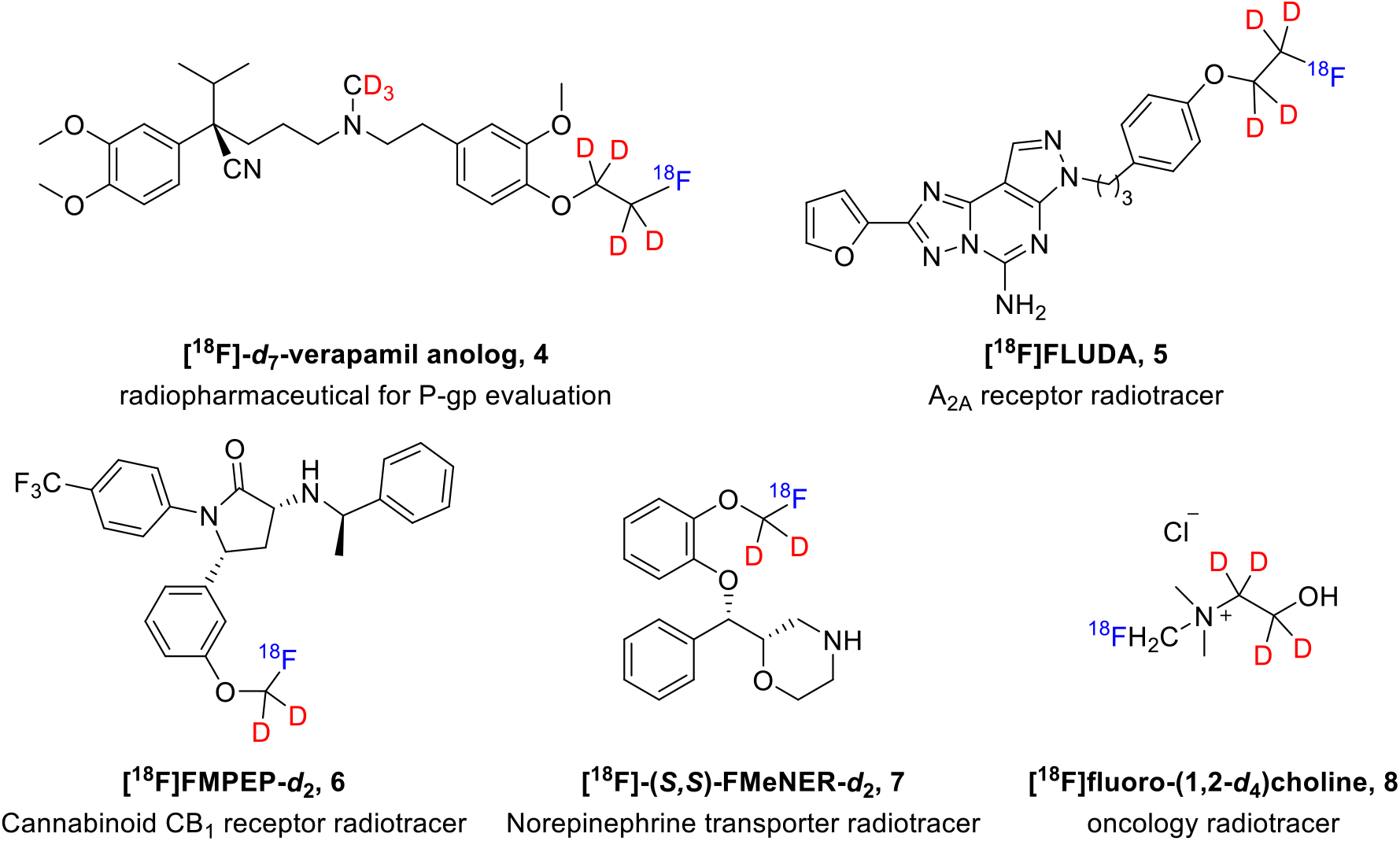
Examples of deuterated PET radiotracers.

Herein we investigate the enzymes responsible for the metabolism of [^18^F]3F4AP, the stability of deuterated derivatives, and the identity of its metabolites to better understand the tracer’s metabolic stability and potentially develop strategies to improve upon it.

## Materials and Methods

### General

All chemicals were ordered from commercial suppliers and used without further purification. 4-aminopyridine-d_6_ (98% deuterium incorporation) was purchased from CDN isotopes. The protium/deuterium exchange reaction was conducted with Biotage^®^ Initiator^+^ microwave synthesizer. The ^1^H, ^19^F and ^13^C NMR spectra were collected on a 300 MHz Bruker spectrometer at 300 MHz, 282 MHz, and 75 MHz, respectively. Copies of the spectra can be found in the supporting information. All ^1^H-NMR data are reported in δ units, parts per million (ppm), and were calibrated relative to the signals for residual chloroform (7.26 ppm) in deuterochloroform (CDCl_3_). All ^13^C-NMR data are reported in ppm relative to CDCl_3_ (77.16 ppm) and were obtained with ^1^H decoupling unless otherwise stated. The following abbreviations or combinations thereof were used to explain the multiplicities: s = singlet, d = doublet, t = triplet, q = quartet, br = broad, m = multiplet. High-resolution mass spectra (HRMS) were recorded on a Thermo Scientific Dionex Ultimate 3000 UHPLC coupled to a Thermo Q Exactive Plus mass spectrometer system using ESI as the ionization approach.

The examination of the relative metabolic rates was carried out with Life Technologies™ Vivid^®^ CYP2E1 screening kit, which includes reaction buffer (2X, 400 mM potassium phosphate pH 8.0), CYP2E1 BACULOSOMES^®^ Plus reagent (consist of recombinant human cytochrome P450, human cytochrome P450 reductase, human cytochrome b_5_), Vivid^®^ regeneration system (100X, 333 mM Glucose-6-phosphate and 30 Units/mL Glucose-6-phosphate dehydrogenases in 100 mM potassium phosphate, pH 8.0), Vivid^®^ NADP^+^ (10mM NADP^+^ in 100 mM potassium phosphate, pH 8.0), Vivid^®^ EOMCC substrate (2H-1-benzopyran-3-carbonitrile,7-(ethoxy-methoxy)-2-oxo-(9Cl)), Vivid^®^ blue fluorescent standard.

The competitive reaction with the Vivid^®^ CYP2E1 screening kit was conducted with Falcon^®^ 384-well Optilux black/clear flat bottom TC-treated micro-test microplates at room temperature with 3 replicates. *Trans*-2-phenylcyclopropylamine hydrochloride was used as a control inhibitor. The fluorescence measurement was performed in kinetic assay mode (reads in 1-minute intervals over 60 min) with Molecular Devices SpectraMax M3 Multi-Mode microplate reader (fluorescence, excitation wavelength = 415 nm, emission wavelength = 460 nm).

The Vivid^®^ CYP2E1 catalyzed nonradioactive and radioactive 3F4AP reaction was conducted in an incubator shaker (BTLab systems BT921) at 37 °C.

The [^18^F]3F4AP was produced in the GE TRACERlab Fx2N synthesizer. Semipreparative HPLC separations were performed on Sykam S1122 Solvent Delivery System HPLC pump with the UV detector at 254 nm with a Waters C18 preparative column (XBridge BEH C18 OBD Prep Column 130 Å, 5 µm, 10 mm × 250 mm).

Other analytical and preparative HPLC analysis was conducted on a Thermo Scientific Dionex Ultimate 3000 UHPLC equipped with below columns as appropriate: Waters XBridge BEH HILIC analytical column (3.5 µm, 4.6 × 150 mm); Waters XBridge BEH C18 analytical column (130 Å, 3.5 µm, 4.6 × 100 mm); SiliChrom HILIC semipreparative column (100 Å, 5µm, 10 × 150 mm).

### Experimental procedures

#### Verifying 3F4AP relative potency towards CYP2E1

In a falcon black/clear 384-well plate, 40 µL of 2.5X (final concentration 15µM) test compounds (4AP, 3F4AP, tranylcypromine) solution in 1X Vivid^®^ CYP2E1 reaction buffer was added to desired wells with three replicates. After 50 µL master pre-mix (2X (40 nM) CYP2E1 BACULOSOMES^®^ and 2X (0.6 Units/mL) Vivid^®^ regeneration system in 1X reaction buffer) was added to each well, the plate was incubated for 10 minutes at room temperature to allow the compounds to interact with the CYP2E1 in the absence of enzyme turnover. At last, reaction was initiated by adding 10µL per well of 10X (100 μM) Vivid^®^ substrate and 10X (300 μM) Vivid^®^ NADP^+^ mixture. Immediately (less than 2 minutes), the plate was transferred into the fluorescent plate reader and fluorescence was monitored over 60 minutes (reads in 1-minute intervals) at 415 nm as excitation wavelength and 460 nm as emission wavelength. The obtained reads were plotted using GraphPad Prism 9.

#### *Determination of the IC*_*50*_ *and* K_i_ *of 4AP and 3F4AP*

A similar Vivid^®^ CYP2E1 assay was conducted as described above. Instead of testing a single concentration of the test compounds (final concentration 15µM), a series of concentrations of 4AP (120mM, 40 mM, 12 mM, 4.0 mM, 1.2 mM, 400 µM, 120 µM, 40 µM, 12 µM, 4.0 µM), 3F4AP (400 µM, 120 µM, 40 µM, 12 µM, 4.0 µM, 1.2 µM, 0.4 µM) and tranylcypromine (40 µM, 12 µM, 4.0 µM, 1.2 µM, 0.4 µM, 0.12 µM, 0.04 µM) were tested with 3 replicates for each concentration. The plate fluorescence was monitored over 60 minutes (reads in 1-minute intervals) at 415 nm as excitation wavelength and 460 nm as emission wavelength. The reads at 60 min (recalculated by the linear trend line equation) of each concentration were used and fitted with GraphPad Prism9 dose-response-inhibition (concentration is log) curve fitting to calculate the IC_50_ values. The corresponding *K*_*i*_ values were calculated by *K*_*i*_ = IC_50_/(1+[S]/*K*_*m*_ (*K*_*m*_ = 22 (Bryan D. Marks and Mary S. Ozers 2002), the Vivid^®^ EOMCC substrate concentration [S] is 10 µM).

#### *Synthesis of 3-fluoro-4-aminopyridine-5-*d_*1*_ (2-*d*_1_*) via H/D exchange in the presence of DCl*

A solution of 3-fluoro-4-aminopyridine (**2**) (56 mg, 0.5 mmol) in D_2_O (1.0 mL) in the presence of DCl (37% in D_2_O, 1.0 mL) was irradiated in a sealed Biotage microwave tube at 170 °C for 12 h (hold-time) using Biotage microwave synthesizer (maximum pressure 12 bar) by moderation of the initial microwave power (400 W). The mixture was cooled in a stream of compressed air and neutralized with aqueous NaOH solution (5 M, 2.5 mL). The solvent (water) was evaporated under reduced pressure and the residue was redissolved with CH_2_Cl_2_ (3 × 2 mL). The combined organic solution was dried over magnesium sulfate, filtered, and evaporated under reduced pressure to give deuterated isotopologue 3-fluoro-4-aminopyridine-5-*d*_1_ (**2-*d***_**1**_) in 77% yield. ^**1**^**H NMR** (300 MHz, CDCl_3_): δ 8.15 (s, 0.95 H), 7.99 (s, 0.97 H), 6.62 (dd, *J* = 7.0, 5.7 Hz, 0.05 H), 4.07 (br, 2 H) ppm. ^**13**^**C NMR** (75 MHz, CDCl_3_): δ 149.21 (d, *J*_*C-F*_ = 246.2 Hz), 146.14, 146.04 (d, *J*_*C-F*_ = 4.8 Hz), 141.44 (d, *J*_*C-F*_ = 10.6 Hz), 136.60 (d, *J*_*C-F*_ = 20.1 Hz), 110.90 (d, *J*_*C-F*_ = 2.0 Hz), 110.63 (td, *J*_*C-D*_ = 24.7 Hz, *J*_*C-F*_ = 1.9 Hz) ppm. ^**19**^**F NMR** (282 MHz, CDCl_3_): δ 152.3 ppm. **ESI-HRMS** (m/z): [M+H]^+^ calc’d for C_5_H_5_D_1_FN_2_^+^: 114.0572; found: 114.0573 with 94.5%; C_5_H_6_FN_2_^+^: 113.0510; found: 113.0510 with 5.5%.

#### *Synthesis of 3-fluoro-4-aminopyridine-2,6-*d_*2*_ (2-*d*_*2*_) *and 3-fluoro-4-aminopyridine-2-*d_*1*_ (2-*d*_*1*_’) *via H/D exchange under neutral condition*

A solution of 3-fluoro-4-aminopyridine (**2**) (56 mg, 0.5 mmol) in D_2_O (2.0 mL) was irradiated in a sealed Biotage microwave tube at 170 °C for 12 h (hold-time) using Biotage microwave synthesizer (maximum pressure 12 bar) by moderation of the initial microwave power (400 W). The mixture was cooled in a stream of compressed air, then acidified with 1.0 mL HCl (1 M in aqueous) and neutralized with aqueous NaOH solution (1 M, 1.0 mL). The solvent (water) was evaporated under reduced pressure and the residue was redissolved with CH_2_Cl_2_ (3 × 2 mL). The combined organic solution was dried over magnesium sulfate, filtered, and evaporated under reduced pressure to give deuterated isotopologues 3-fluoro-4-aminopyridine-2,6-*d*_2_ (**2-*d***_**2**_) and 3-fluoro-4-aminopyridine-2-*d*_1_ (**2-*d***_**1**_**’**) mixture in 84%. ^**1**^**H NMR** (300 MHz, CDCl_3_): δ 8.13 (s, 0.02 H), 7.97 (d, *J* = 4.81 Hz, 0.52 H), 6.61 (m, 0.94 H), 4.59 (br, 2 H) ppm. ^**13**^**C NMR** (75 MHz, CDCl_3_): δ 149.04 (d, *J*_*C-F*_ = 246.1 Hz), 145.99 (td, *J*_*C-D*_ = 3.1, *J*_*C-F*_ = 1.3 Hz), 145.66 (tdt, *J*_*C-D*_ = 27.3, 1.7 Hz, *J*_*C-F*_ = 4.7 Hz), 141.63 (d, *J*_*C-F*_ = 10.5 Hz), 136.17 (td, *J*_*C-D*_ = 27.4 Hz, *J*_*C-F*_ = 20.2 Hz), 110.87 (d, *J*_*C-F*_ = 2.0 Hz), 110.74 (br) ppm. ^**19**^**F NMR** (282 MHz, CDCl_3_): δ 152.41, 152.44 ppm. **ESI-HRMS** (m/z): [M+H]^+^ calc’d for C_5_H_4_D_2_FN_2_^+^: 115.0635, found: 115.0635, 46.8%; C_5_H_5_DFN_2_^+^: 114.0572, found: 114.0573, 52.0%; C_5_H_6_FN_2_ ^+^: 113.0510, found: 113.0511, 1.2%.

#### *Synthesis of 3-fluoro-4-aminopyridine-2,5,6-*d_*3*_ (2-*d*_3_*) via H/D exchange under neutral condition*

A solution of 3-fluoro-4-aminopyridine-5-*d*_1_ (**2-*d***_**1**_) (18 mg, 0.2 mmol) in D_2_O (1.0 mL) was irradiated in a sealed Biotage Microwave tube at 190 °C for 8 h (hold-time) using Biotage microwave synthesizer (maximum pressure 12 bar) by moderation of the initial microwave power (400 W). The mixture was cooled in a stream of compressed air, then acidified with 1.0 mL HCl (1 M in aqueous) and neutralized with aqueous NaOH solution (1 M, 1.0 mL). The solvent (water) was evaporated under reduced pressure and the residue was redissolved with CH_2_Cl_2_ (3 × 2 mL). The combined organic solution was dried over magnesium sulfate, filtered, and evaporated under reduced pressure to give deuterated isotopologue 3-fluoro-4-aminopyridine-2,5,6-*d*_3_ (**2-*d***_**3**_) in 76% yield. ^**1**^**H NMR** (300 MHz, CDCl_3_): δ 8.12 (s, 0.02 H), 7.97 (s, 0.07 H), 6.61 (d, *J* = 7.6, 0.03 H), 4.59 (br, 2 H) ppm. ^**13**^**C NMR** (75 MHz, CDCl_3_): δ 149.04 (d, *J*_*C-F*_ = 246.1 Hz), 145.94, 145.62 (tm, *J*_*C-D*_ = 27.0 Hz), 141.60 (d, *J*_*C-F*_ = 10.6 Hz), 136.15 (td, *J*_*C-D*_ = 27.0 Hz, *J*_*C-F*_ = 20.0 Hz), 110.46 (td, *J*_*C-D*_ = 24.7 Hz) ppm. ^**19**^**F NMR** (282 MHz, CDCl_3_): δ 152.5 ppm. **ESI-HRMS** (m/z): [M+H]^+^ calc’d for C_5_H_3_D_3_FN_2_^+^: 116.0698, found: 116.0696, 87.1%; C_5_H_4_D_2_FN_2_^+^: 115.0635, found: 115.0634, 13.1%.

#### In vitro *evaluation of deuterated 3F4AP analogues with the human enzyme*

In a falcon black/clear 384-well plate, 40 µL of 2.5X test compounds (4AP, 4AP-*d*_6_, 3F4AP, 3F4AP-*d*_x_, tranylcypromine) solution in 1X Vivid^®^ CYP2E1 reaction buffer was added to desired wells with three replicates. After 50 µL master pre-mix (2X (40 nM) CYP2E1 BACULOSOMES^®^ and 2X (0.6 Units/mL) Vivid^®^ regeneration system in 1X reaction buffer) was added to each well, the plate was incubated for 10 minutes at room temperature to allow the compounds to interact with the CYP2E1 in the absence of enzyme turnover. Reaction was initiated by adding 10µL per well of 10X (100 µM) Vivid^®^ substrate and 10X (300 µM) Vivid^®^ NADP^+^ mixture. Immediately (less than 2 minutes), plate was transferred into the fluorescent plate reader and fluorescence was monitored over 60 minutes (reads in 1-minute intervals) at 415 nm as excitation wavelength and 460 nm as emission wavelength. The obtained reads were plotted using GraphPad Prism 9.

#### In vitro *identification of metabolites with the human enzyme by high resolution mass spectrometry*

The reactions were conducted with the Vivid^®^ CYP2E1 enzyme, NADP^+^, and NADPH regeneration system. In an Eppendorf vial, 40 µL of 2.5X deuterated substrates (3F4AP-*d*_x_, 125 µM) solution in 1X Vivid^®^ CYP2E1 reaction buffer was added. After 50 µL master pre-mix (2X (200 nM) CYP2E1 BACULOSOMES^®^ and 2X (3 Units/mL) Vivid^®^ regeneration system in 1X reaction buffer) was added to each reaction vial, the reaction mixture was incubated for 10 minutes at 37 ºC to allow the compounds to interact with the CYP2E1 in the absence of enzyme turnover. Reaction was initiated by adding 10 µL per well of 10X Vivid^®^ NADP^+^ mixture (1.5 mM). Immediately (less than 2 minutes), the vial was transferred into a 37 ºC shaker and allowed to react for 20 hours. Reaction mixture was then diluted to 5 folds with ultrapure water and analyzed with high-resolution mass spectrometry.

#### Synthesis of 3-fluoro-4-aminopyridine 1-oxide (3F4AP N-oxide, 9a)

A suspension of 3-fluoro-4-nitropyridine 1-oxide (**10**) (50 mg, 0.32 mmol) and palladium on carbon (10%, 5.0 mg) in ethanol (2.0 mL) was bubbled with Argon gas for 10 minutes. The reaction mixture was cooled down to 0 °C. A hydrogen balloon was loaded and bubbled for another 5 minutes before removing the pressure balance needle. The reaction mixture was kept under stirring and allowed to warm up to room temperature slowly. The reaction was monitored by TLC and analytical HPLC with a C18 analytical column (mobile phase A: 10 mM NH_4_HCO_3_ aqueous solution pH 8.0; mobile phase B: acetonitrile; gradient method: 0-5 min: 3% B, 5-8 min: 3% to 90% B, 8-11 min: 90% B, 11-12 min: 90% to 3% B, 12-15 min: 3% B; t_R_ of **10** is 8.0 min). Pd/C was filtered out after **10** was consumed completely. The filtrate was concentrated under reduced pressure, and the residue was purified with prep-HPLC (HILIC semipreparative column, 10 mM NH_4_HCO_3_ aqueous solution pH 8.0/acetonitrile = 10/90) to afford **3c** as an off-white solid in an 84% yield. ^**1**^**H NMR** (300 MHz, MeOD) δ 8.27 – 8.05 (m, 1H), 7.84 (d, *J* = 7.1 Hz, 1H), 6.82 (t, *J* = 8.5 Hz, 1H) ppm. ^**13**^**C NMR** (75 MHz, MeOD) δ 148.54 (d, *J* = 245.4 Hz), 142.29 (d, *J* = 12.0 Hz), 137.30, 129.28 (d, *J* = 33.0 Hz), 111.40 (d, *J* = 6.5 Hz) ppm. ^**19**^**F NMR** (282 MHz, MeOD) δ −146.00 (dd, *J* = 9.9, 6.3 Hz) ppm. **ESI-HRMS** (m/z): [M+H]^+^ calc’d for C_5_H_5_FN_2_O^+^: 129.0459, found: 129.0457.

#### Preparation of [^18^F]3F4AP sample for Vivid^®^ CYP2E1 catalyzed reaction

[^18^F]3F4AP was prepared with GE Fx2N according to the reported procedure (Basuli et al. 2018). The obtained [^18^F]3F4AP fraction (radiopurity > 99%, molar activity = 3.8 ± 0.5 Ci/μmol, n = 2, in 95% 20 mM sodium phosphate buffer, pH 8.0, 5% EtOH solution) was concentrated at 10% of volume to reduce EtOH amount as much as possible. The resulted solution was used directly for further reaction.

#### *Verifying the presence of 3F4AP N-oxide metabolite with the human enzyme* in vitro *by radio HPLC*

The reactions were conducted with Vivid^®^ CYP2E1 enzyme, NADP^+^, and NADPH regeneration system. In a 1.5 mL Eppendorf vial, 10 µL of [^18^F]3F4AP solution above was diluted with 30 µL of 1X Vivid^®^ CYP2E1 reaction buffer. After 50 µL master pre-mix (2X, 200 nM CYP2E1 BACULOSOMES^®^ and 2X Vivid^®^ regeneration system 3 Units/mL in 1X reaction buffer) was added, the reaction mixture was incubated for 10 minutes at 37 ºC to allow [^18^F]3F4AP to interact with the CYP2E1 in the absence of enzyme turnover. Reaction was initiated by adding 10µL of 10X Vivid^®^ NADP^+^ mixture (1.5 mM). Immediately, reaction vial was transferred into a 37 ºC shaker and allowed to react for 2 hours. The reaction mixture was then filtrated with Amicon Ultra-0.5 centrifugal filter (pore size, 10 kDa NMWCO). The filtrate was analyzed with HPLC through a HILIC analytical column (mobile phase: 10 mM NH_4_HCO_3_ aqueous solution pH 8.0/acetonitrile 10/90). The identification of the formation of [^18^F]**9a** (t_R_ ∼ 6.0 min in radio chromatogram) was confirmed by co-injection with **9a** (t_R_ ∼ 5.9 min in UV chromatogram).

## Results

In the following sections, we provide evidence that CYP2E1 is the predominant enzyme responsible for 3F4AP metabolism. We also describe the chemical synthesis of several deuterated isotopologues of 3F4AP and compare their relative rate towards CYP2E1 mediated oxidation. Finally, through a combination of HRMS and radio-HPLC we identify the primary metabolites.

### Cytochrome P450 CYP2E1 metabolizes 3F4AP

To examine that CYP2E1 is responsible for 3F4AP metabolism, we compared the reaction rate of 4AP, 3F4AP and tranylcypromine (positive inhibition) using the Life Technologies™ Vivid^®^ CYP2E1 assay. This kit relies on the detection of fluorescence emitted by 3-cyano-7-hydroxy-coumarin, the product from the *O*-dealkylation reaction of a specific CYP2E1 substrate (2H-1-benzopyran-3-carbonitrile,7-(ethoxy-methoxy)-2-oxo-(9Cl)) by this enzyme in the absence and presence of substrate competitors (Bryan D. Marks and Mary S. Ozers 2002). Based on this kit’s principle, the blank experiments (without any competitors) result in the highest fluorescence values while increasing amounts of competitors decrease the fluorogenic emission.

As it can be seen in Figure 3A, tranylcypromine was very effective at slowing down the formation of the fluorescent product, indicating that it is an excellent inhibitor of the CYP2E1 (maroon *vs*. blue lines). In comparison, 4AP showed a moderate reduction in the reaction rate (cyan *vs*. blue lines), indicating that it is a poorer substrate. 3F4AP, on the other hand, showed a very robust reduction in the reaction rate (orange line), indicating that it is also a very good substrate of the enzyme.

**Figure 3.**
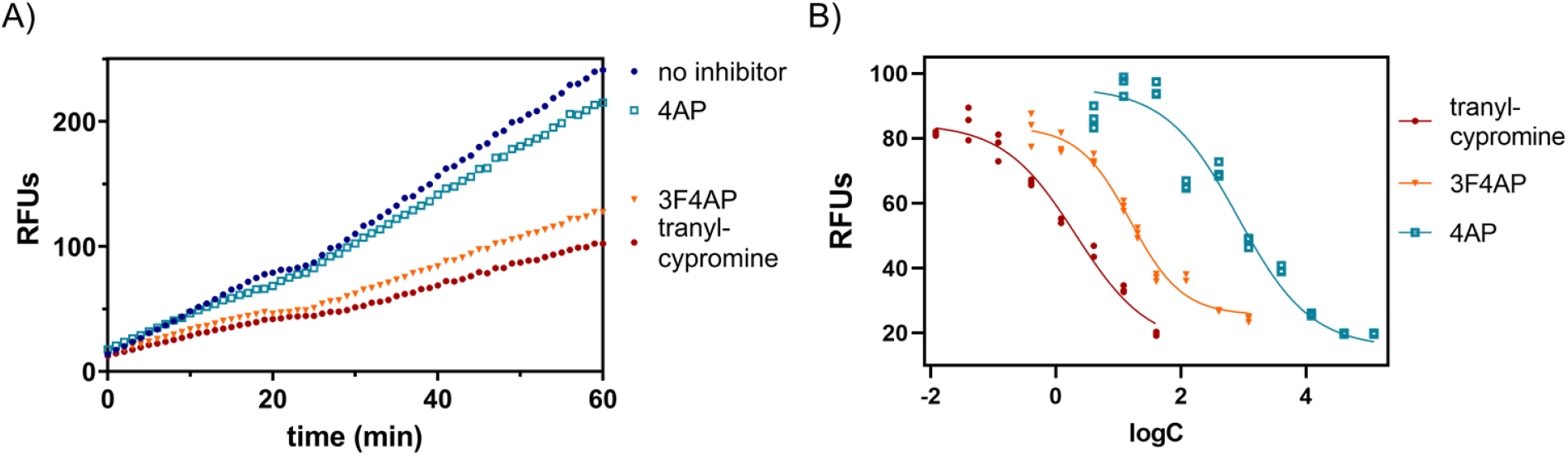
Evaluation of CYP2E1 enzyme-catalyzed oxidation of 3F4AP. A) Relative fluorescence-time curves based on the 60 minutes kinetic measurement of 4AP, 3F4AP, and tranylcypromine as positive control for inhibition; B) IC_50_ fit curves of 4AP, 3F4AP, and tranylcypromine.

In order to quantify these differences, we measured the reaction rate in the presence of different concentrations of 4AP, 3F4AP, and tranylcypromine. Plotting the relative fluorescence units at 60 min (RFUs) against the logarithm of the concentrations resulted in the sigmoidal curves, which could be fitted to calculate the IC_50_ and *K*_i_ (Fig. 3B). As shown in **Table 1**, the calculated IC_50_ and *K*_*i*_ of tranylcypromine were largely consistent with previously published values (Bryan D. Marks and Mary S. Ozers 2002). In comparison, 4AP showed a ∼400-fold higher IC_50_ and *K*_i_, confirming that it is a weaker substrate. 3F4AP, on the other hand, showed an IC_50_ and *K*_i_ around 8-fold higher than tranylcypromine which confirms that 3F4AP is an excellent substrate of CYP2E1. For reference acetaminophen, a known substrate of CYP2E1, has much higher IC_50_ and *K*_i_ values than 4AP, confirming that both 4AP and 3F4AP are suitable CYP2E1 substrates, although 3F4AP is a much better one. This explains why in humans 85% of unmetabolized 4AP is recovered 24h after a single oral or intravenous administration (Uges et al. 1982; Evenhuism et al. 1981), whereas less than 50% of unmetabolized [^18^F]3F4AP remains in circulation 1h post intravenous administration.

**Table 1.** IC_50_ and *K*_*i*_ values for CYP2E1 substrates.

| compounds | Measured values |  | Published values |  |
| --- | --- | --- | --- | --- |
|  | IC<sub>50</sub>(μM) | K<sub>i</sub>(μM) | IC<sub>50</sub>(μM) | K<sub>i</sub>(μM) |
| tranylcypromine | 2.06 | 1.6 | 0.96 | 0.6 |
| 4-aminopyridine | 810 | 550 | -- | -- |
| 3-fluoro-4-aminopyridine | 15.4 | 10.6 | -- | -- |
| Acetaminophen | -- | -- | 1700 | 1100 |

### Synthesis of deuterated 3F4AP isotopologues and their metabolic stability

Given that 3F4AP is an excellent substrate of CYP2E1, we sought to test the hypothesis that deuterated forms of 3F4AP would be more stable against CYP2E1 oxidation. Thus, we prepared the deuterated 3F4AP isotopologues and tested their relative metabolic stability using the Vivid^®^ CYP2E1 assay.

Attempts to fluorinate 4AP-*d*_6_ with reactive electrophilic fluorination reagent SelectFluor in water or a combined water/chloroform solvent system at 80 °C did not generate deuterated 3F4AP in useful yields (<5% conversion), prompting us to explore the direct protium/deuterium exchange of 3F4AP. We decided to test H/D exchange under microwave irradiation as it requires lower temperature and shorter reaction time compared to other methods (Bagley et al. 2016; Esaki et al. 2006; C 1989). As shown in Figure 4, reaction of 3F4AP in D_2_O under microwave irradiation at 170 °C gave 2,6-*d*_2_-3F4AP with D incorporation of 98% at C-2 and 48% at C-6 after 12 hours. On the other hand, the reaction in the presence of DCl at 170 °C produced 5-*d*_1_-3F4AP with D incorporation 94% at C-5. The regioselectivity of this acid-mediated deuteration, preferably at *β*-position, can be explained by an aromatic electrophilic substitution mechanism. To synthesize a fully deuterated 3F4AP, a sequential reaction with both conditions was conducted, resulting in 2,5,6-*d*_3_-3F4AP with D incorporation of 97%, 98%, 93% at C-5, 2, 6, respectively. Moreover, these straightforward protium/deuterium exchange reactions gave a 76-84% yield range.

**Figure 4.**
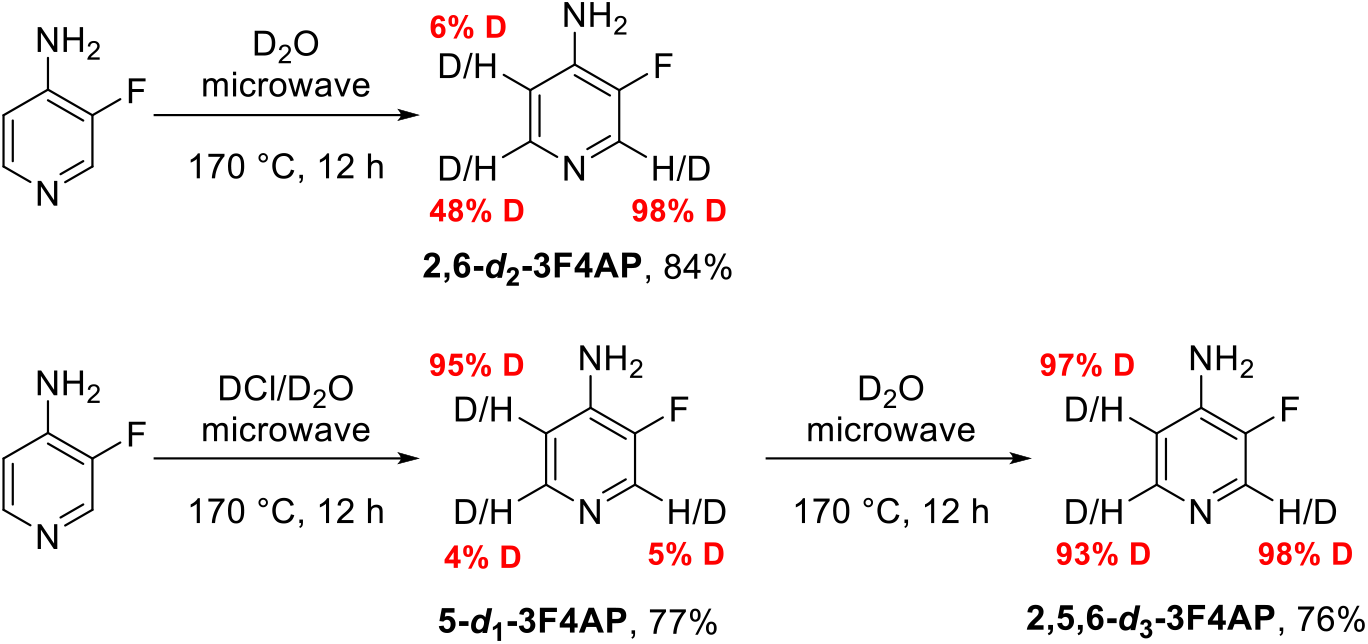
Deuteration of 3F4AP under microwave irradiation.

Then, we evaluated the kinetic isotope effect of the prepared deuterated 3F4AP isotopologues as well as the commercially available 4AP-d_6_ using the Vivid^®^ CYP2E1 assay. As shown in Figure 5, the fluorescence response curves of deuterated isotopologues are almost equivalent to their corresponding non-deuterated compounds indicating no deuterium kinetic isotope effect (KIEs < 1.1 for all the deuterated isotopologues).

**Figure 5.**
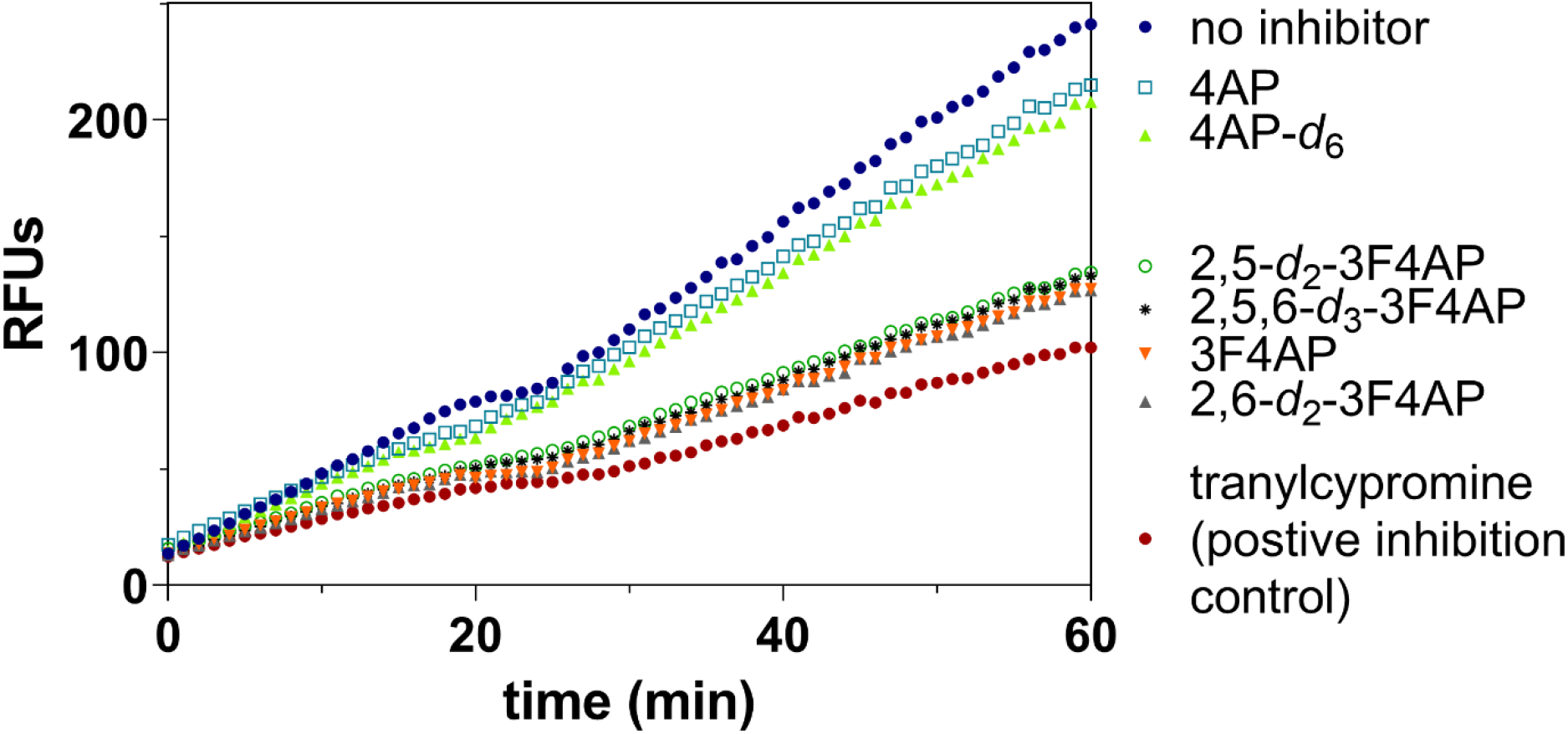
Evaluation of kinetic isotope effect in the Vivid^®^ CYP2E1 enzyme-catalyzed oxidation. Relative fluorescence-time curves based on the 60 minutes kinetic measurement of 4AP, 4AP-*d*_6_, 3F4AP, and three deuterated 3F4AP-*d*_x_; tranylcypromine is known as a good CYP2E1 inhibitor.

### Identification of the oxidation metabolites of 3F4AP

The identification of metabolites of PET tracers can shed light on their clearance mechanism and enable the evaluation of whether any radiometabolites are brain penetrant. Having prepared the deuterated isotopologues we can exploit the difference in mass associated with the loss of a deuterium or hydrogen upon reaction with CYP2E1 to identify the substrate reaction site.

A series of reactions using recombinant CYP2E1 enzyme and the different deuterated 3F4AP substrates (i.e. 2,5,6-*d*_3_-3F4AP, 5-*d*_1_-3F4AP, and 2,6-*d*_2_-3F4AP) were performed and analyzed by high resolution mass spectrometry (HRMS). Figure 6 shows the different reactants and all the possible products with the mass change listed underneath. Oxidation at a hydrogen position results in a mass gain of 16 Da (−1 Da from the loss of one H plus 17 Da from addition of one OH) whereas oxidation at a deuterium position results in a mass gain of 15 Da (−2 Da from the loss of one D plus 17 Da from addition of one OH). Oxidation at the pyridine nitrogen to form the N-oxide product also results in a gain of 16 Da from addition of one oxygen. Since the reaction is performed in aqueous buffer, we assumed that the addition of -OD (plus 17 Da) does not proceed. In reaction A, only the +16 Da was detected, indicating that oxidation at C-2 and C-6 position does not occur. In reactions B and C, we detected both the +15 Da and +16 Da products suggesting that oxidation occurs both at the C-5 and N-1 position.

**Figure 6.**
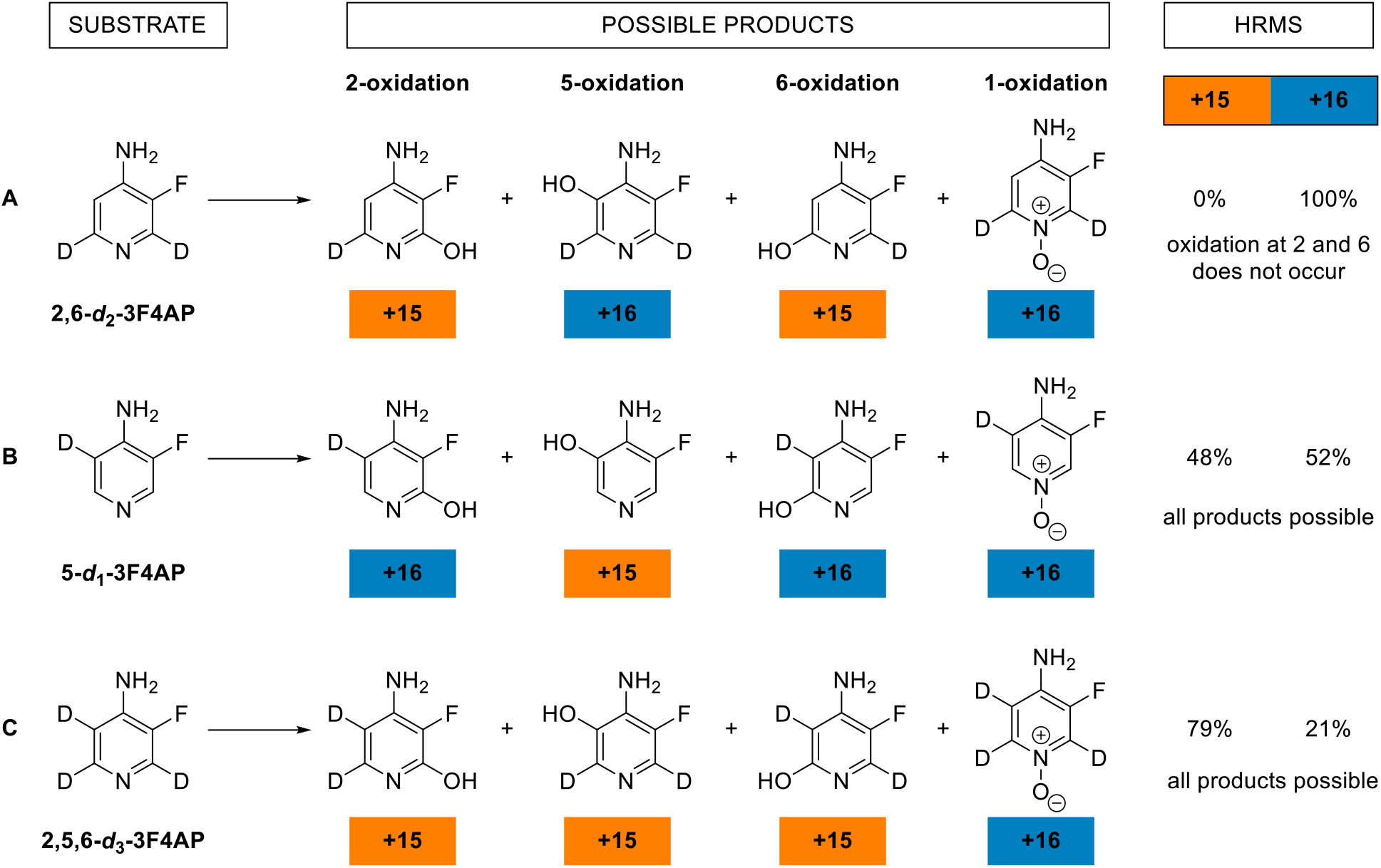
Identification of oxidation metabolites of 3F4AP via HRMS. Vivid^®^ CYP2E1 enzyme-catalyzed oxidation reactions with various deuterated 3F4AP substrates were performed to identify the resulting metabolites via HRMS detection of oxidation at a hydrogen (+16), a deuterium (+15), or the pyridine nitrogen position (+16). See reaction conditions in the methods.

To identify the metabolites produced we performed the CYP2E1 reaction using radioactive [^18^F]3F4AP and looked for the presence of [^18^F]3F4AP N-oxide and 5-hydroxy-[^18^F]3F4AP using radio-HPLC (Fig. 7A). The 3F4AP N-oxide (**9a**) reference standard was prepared *via* Pd/C catalyzed partial reduction of 3-fluoro-4-nitropyridine N-oxide **10** in ethanol under 1 atm of H_2_ (Fig. 7B) and [^18^F]3F4AP was prepared according to the previously reported procedure (Basuli et al. 2018). With these compounds in hand, we performed the CYP2E1 enzymatic reaction on [^18^F]3F4AP and confirmed the formation of [^18^F]3F4AP N-oxide (Fig. 7D, t_R_ ∼ 6.0 min) by co-injection with the prepared standard. In addition, we observed a radiometabolite (t_r_ = 3.7 min), which based on our prior HRMS analysis, can be assigned as 5-hydroxy-3F4AP ([^18^F]**9b)**. The yields of individual metabolites were determined by the area under the curve (AUC) of the corresponding peak. Tracking the reaction at 5, 30, 60, 120, and 240 min, showed that 5-hydroxy-3F4AP ([^18^F]**9b)** is about twice as abundant as [^18^F]3F4AP N-oxide [^18^F]**9a** (Fig. 7E) confirming that the oxidation at the 5 position is more favorable.

**Figure 7.**
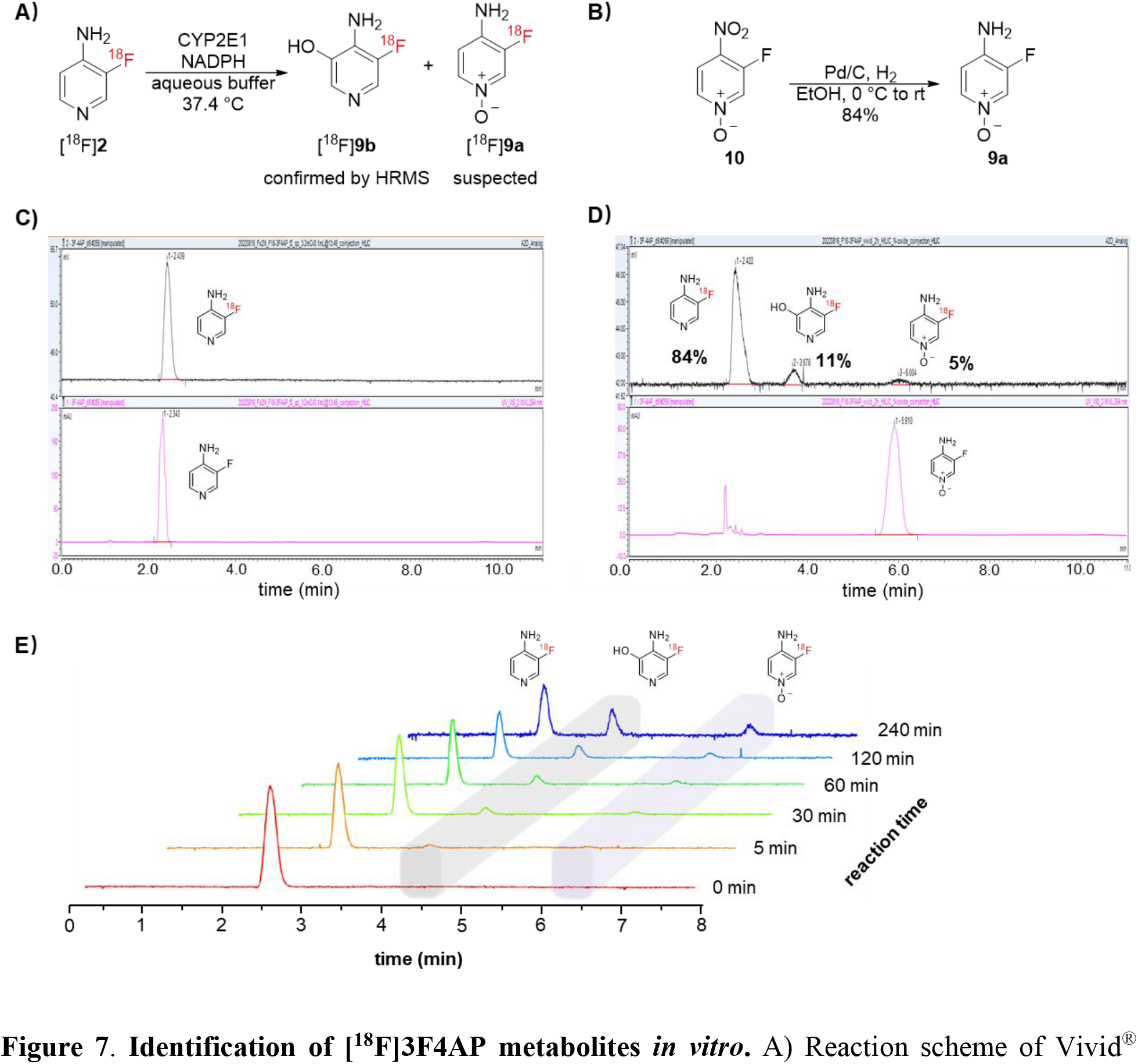
Identification of [^18^F]3F4AP metabolites *in vitro*. A) Reaction scheme of Vivid^®^ CYP2E1 enzyme-catalyzed oxidation with [^18^F]3F4AP as substrate. B) Reaction scheme of 3F4AP N-oxide preparation. Analytical HPLC chromatograms (radio (top) and 254 nm UV (bottom)) of C) [^18^F]3F4AP starting material plus 3F4AP reference standard and D) Vivid^®^ CYP2E1 enzyme-catalyzed oxidation reaction crude (at 240 min) plus 3F4AP N-oxide reference standard. E) Time course of the CYP2E1 enzyme-catalyzed oxidation reaction showing the formation of metabolites at 0, 5, 30, 60, 120 and 240 min.

## Discussion

Understanding the metabolic stability of PET tracers is paramount to its application in humans. Tracers that are metabolized too quickly (in seconds or minutes) may not reach a sufficient concentration in the target, whereas tracers in which the metabolites are present in the target organ may contribute background signal. Furthermore, in order to accurately model the kinetics of a tracer it is necessary to know the fraction of radioactivity corresponding to the parent tracer in circulation. Since metabolism may vary across human subjects understanding the nature of the metabolites and the enzymes responsible for their occurrence can be useful to design strategies to minimize metabolism, develop methods to measure metabolism and understand whether metabolites are present in a target organ such as the brain.

Motivated by recent findings that [^18^F]3F4AP is less stable in awake humans than in anesthetized monkeys and rodents, we investigated the *in vitro* metabolism of 3F4AP and demonstrated that 3F4AP is an excellent substrate of CYP2E1, pointing towards the likely responsibility of this enzyme for the metabolism of 3F4AP *in vivo*. This conclusion is also supported by prior reports indicating that 4AP is primarily metabolized by CYP2E1 and by our experiments with isoflurane-anesthetized and awake mice indicating that isoflurane (a substrate of CYP2E1) can inhibit the metabolism of [^18^F]3F4AP *in vivo*.

In addition, we generated three deuterated 3F4AP isotopologues of 3F4AP by direct protium/deuterium exchange under microwave irradiation with high deuteration incorporation and good yields. These deuterated isotopologues allowed us to probe their stability and identify the metabolites of 3F4AP. In terms of stability, the deuterated isotopologues were not more stable towards oxidation than 3F4AP. In hindsight, this is reasonable given that the deuterium isotope effect is known to occur in aliphatic protons where the C-H bond breaking is the rate-limiting step, whereas in aromatic protons, this is not usually the rate limiting step. This finding further establishes the understanding of when deuteration can be a useful strategy to decrease metabolism.

The deuterated 3F4AP isotopologues were instrumental in identifying the metabolites of 3F4AP as the 5- and 1-oxidation products. Using HRMS, we were able to confirm the formation of 5-hydroxy-3F4AP and 3F4AP N-oxide and exclude the formation of 2- and 6-hydroxy-3F4AP. Additional radio-HPLC experiments using [^18^F]3F4AP confirmed that 5-hydroxy-3F4AP and 3F4AP N-oxide are formed with the former being more abundant.

The identification of the metabolites is valuable information as it allows assessment of whether the radiometabolites of [^18^F]3F4AP can get into the brain. Based on the predicted logP of the identified 3F4AP metabolites (5-hydroxy-3F4AP: −0.34, 3F4AP N-oxide: −1.79), these are not likely brain penetrant, which will facilitate the quantification and interpretation of the PET imaging results with this tracer. This information may also be useful in designing novel PET tracers for detecting demyelination with higher metabolic stability.

## Abbreviations

4AP: 4-aminopyridine
AUC: area under curve
CYP2E1: cytochrome P450 family 2 subfamily E member 1
D_2_O: deuterium oxide
DCl: deuterium chloride
3F4AP: 3-fluoro-4-aminopyridine
HPLC: high-performance liquid chromatography
HRMS: high resolution mass spectrometry
IC_50_: half maximal inhibitory concentration
*K*_*i*_: inhibition constant
KIE: kinetic isotope effect
MS: multiple sclerosis
3OH4AP: 3-hydroxy-4-aminopyridine
PET: positron emission tomography
RFUs: relative fluorescence units.

## Section

Metabolism, Transport, and Pharmacogenomics

## Acknowledgement

The authors thank David F. Lee, Jr., Dr. John A. Correia and Dr. Hamid Sabet for providing the fluorine-18 for the radiotracer synthesis. We thank Jennifer X. Wang for the high-resolution mass spectra measurement and data analysis. We also thank Dr. Marc M. Normandin and Dr. Nicolas J. Guehl for valuable discussion.

## Authorship Contributions

Participated in research design: Sun, Ramos-Torres, Brugarolas

Conducted experiments: Sun, Ramos-Torres

Contributed new reagents or analytic tools: Sun

Performed data analysis: Sun

Wrote or contributed to the writing of the manuscript: Sun, Ramos-Torres, Brugarolas

This study was supported by National Institute of Neurological Disorders and Strokes [Grant R01NS114066].

