## Supplemental information for "Metabolic Stability of the Demyelination PET Tracer [^18^F]3F4AP and Identification of its Metabolites"

### Supporting Information
# of
### Deuterium isotope effects on the stability of the demyelination PET tracer 3F4AP

Yang Sun PhD, Karla Ramos-Torres PhD, and Pedro Brugarolas PhD

Gordon Center for Medical Imaging, Massachusetts General Hospital and Harvard Medical School,

Boston 02114, MA, USA

**Content**

**NMR spectra of 2-$d_1$, 2-$d_2$, 2-$d_3$, and 9a.**

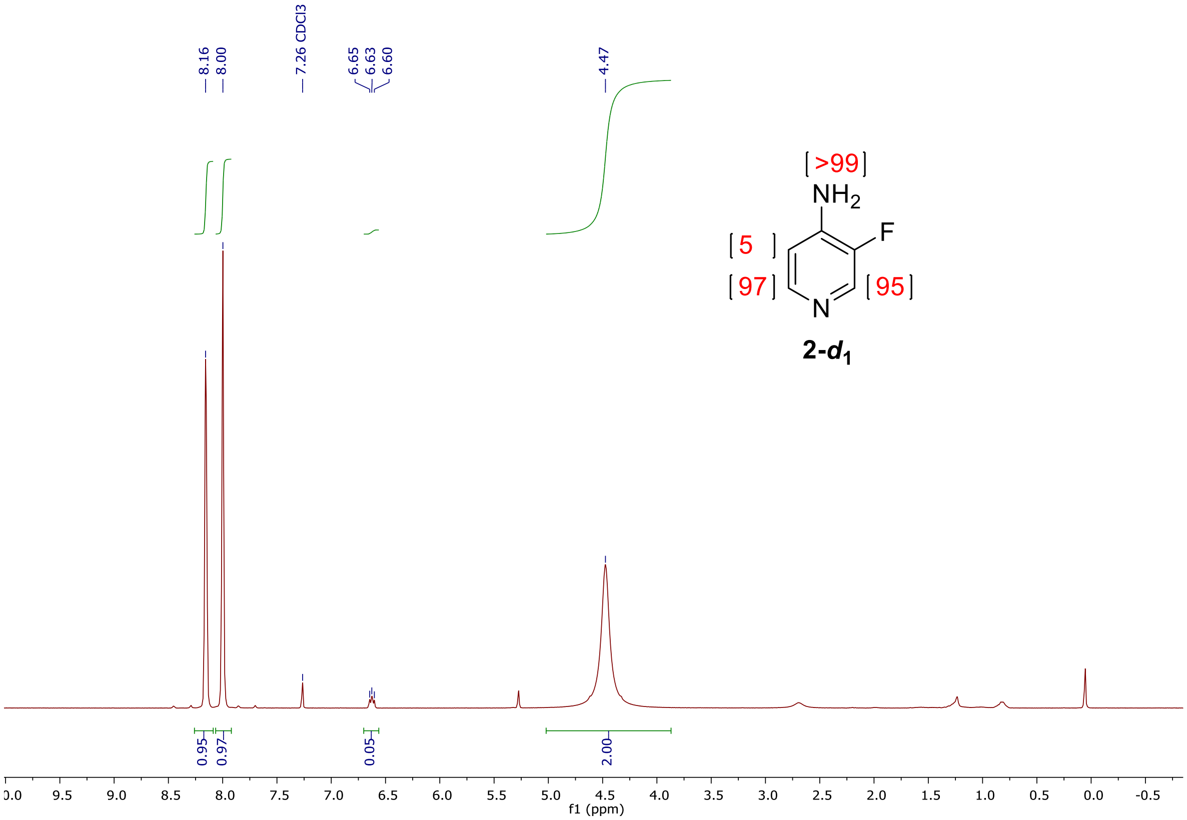


{[1]H NMR} spectrum of 3-fluoro-4-aminopyridine-5-*d*1 (**2-*d*1**).

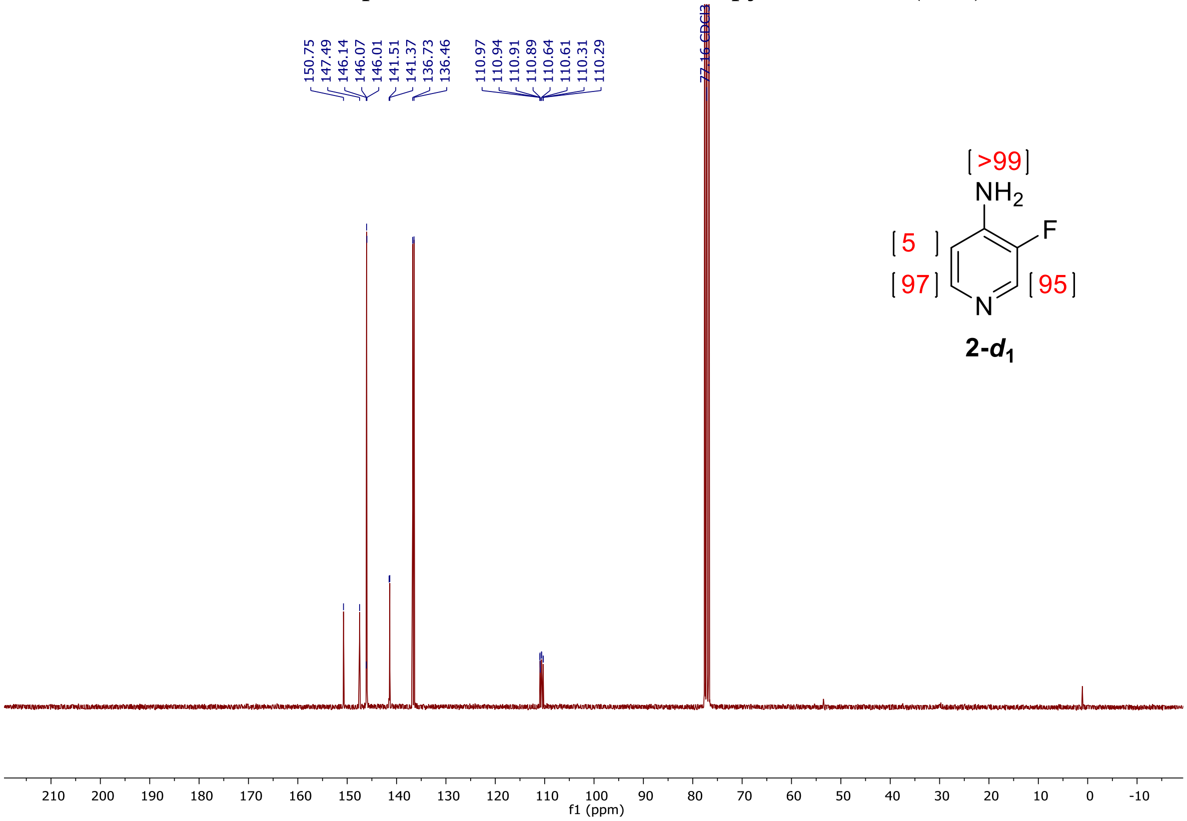


{[13]C NMR} spectrum of 3-fluoro-4-aminopyridine-5-*d*1 (**2-*d*1**).

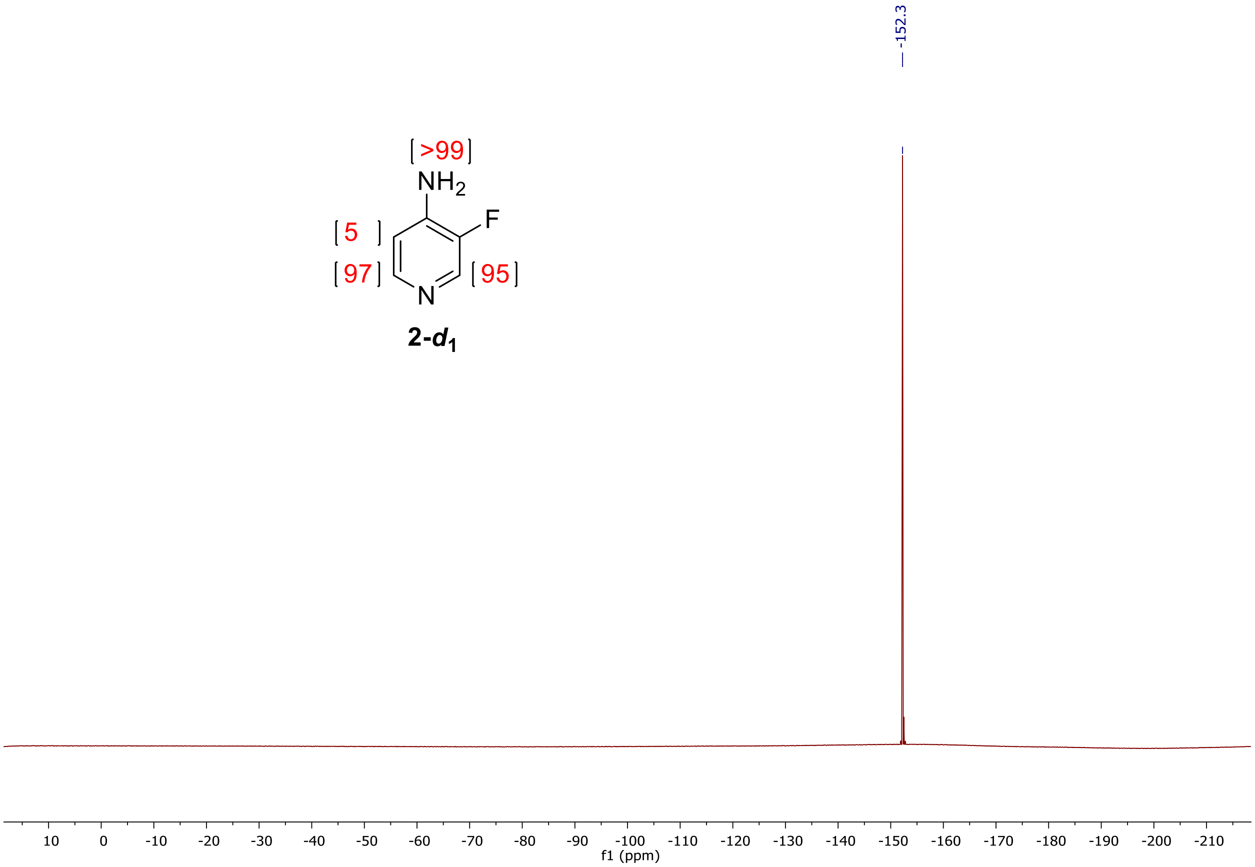


{$^{19}$F NMR} spectrum of 3-fluoro-4-aminopyridine-5-$d_1$ (**2-$d_1$**).

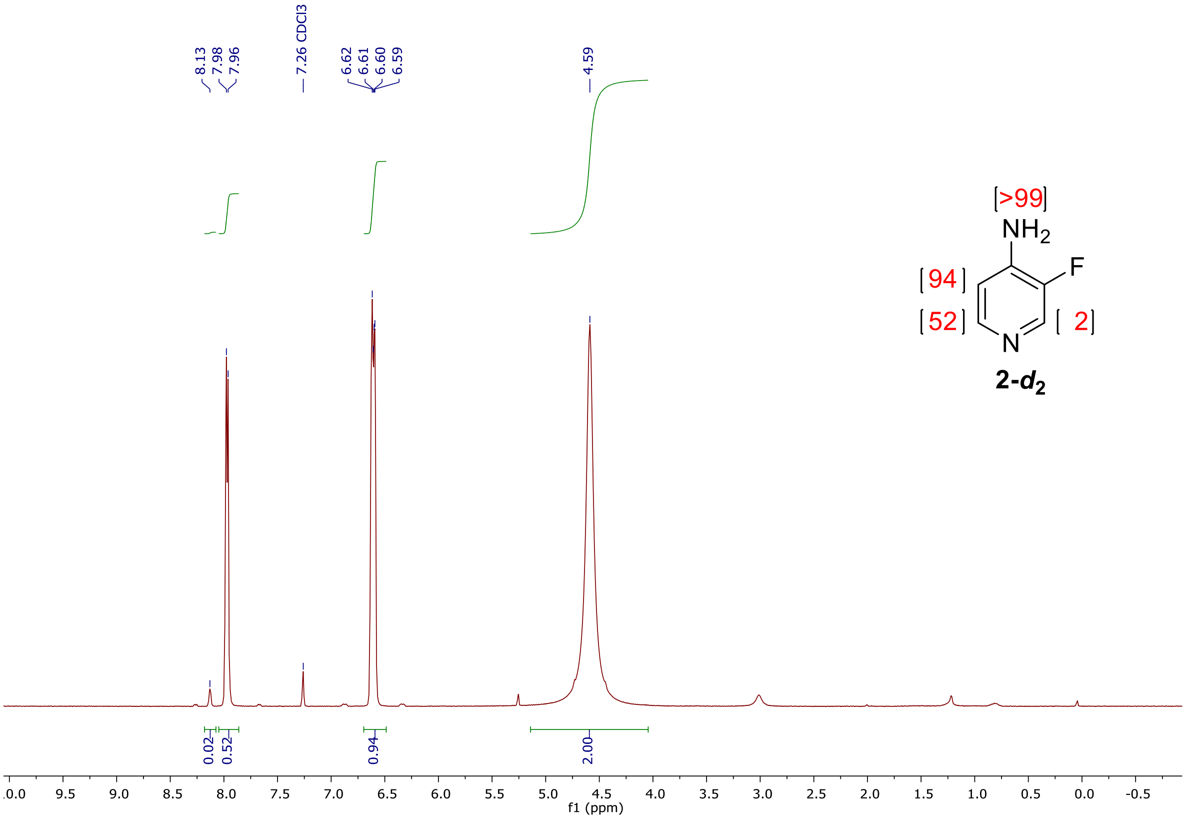


{$^{1}$H NMR} spectrum of 3-fluoro-4-aminopyridine-2,6-$d_2$ (**2-$d_2$**).

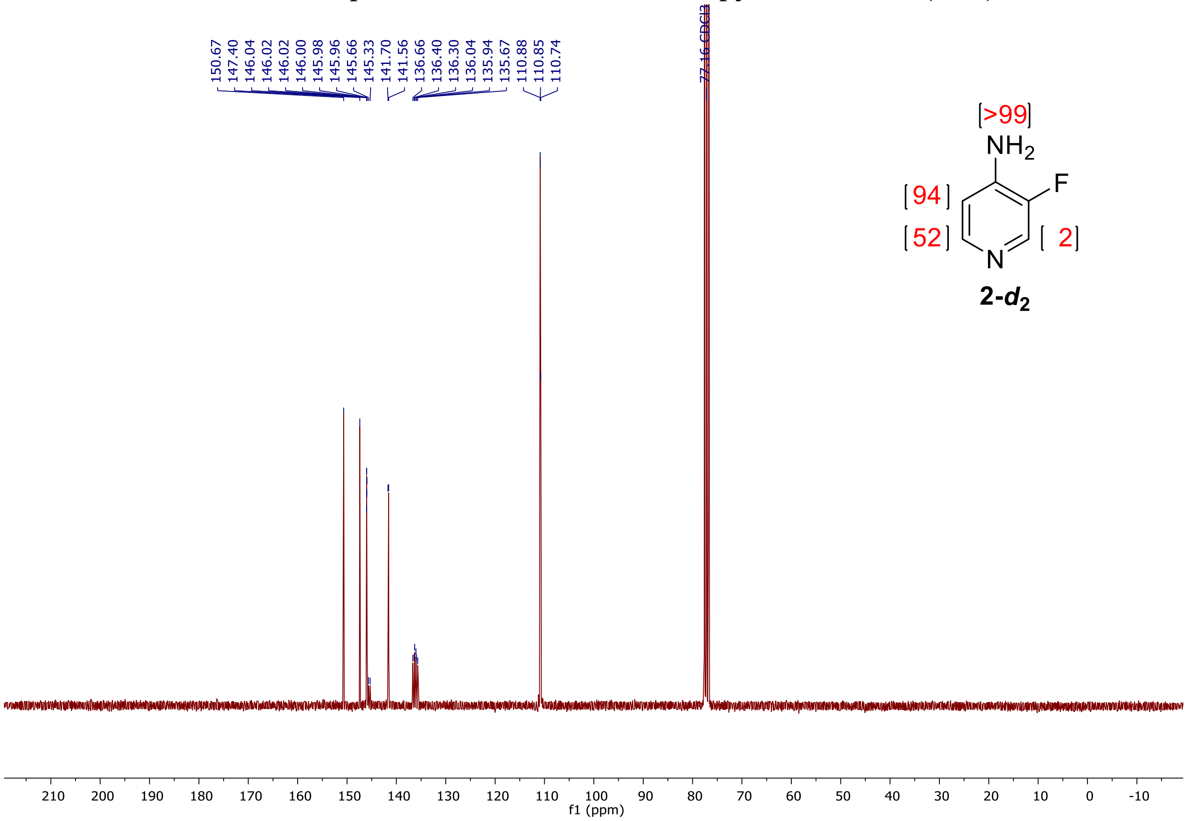


{$^{13}$C NMR} spectrum of 3-fluoro-4-aminopyridine-2,6-$d_2$ (**2-$d_2$**).

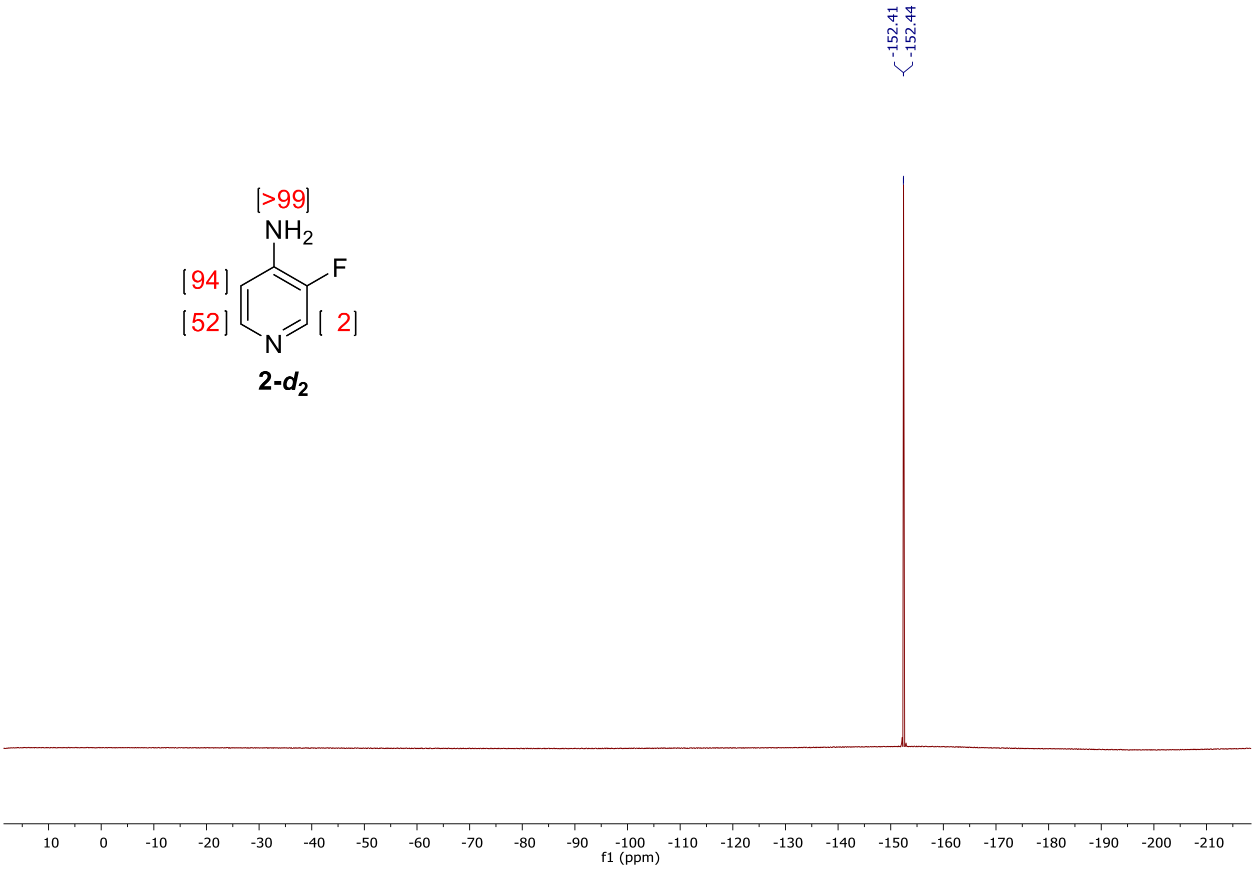


{$^{19}$F NMR} spectrum of 3-fluoro-4-aminopyridine-2,6-*d*$_2$ (**2-*d*$_2$**).

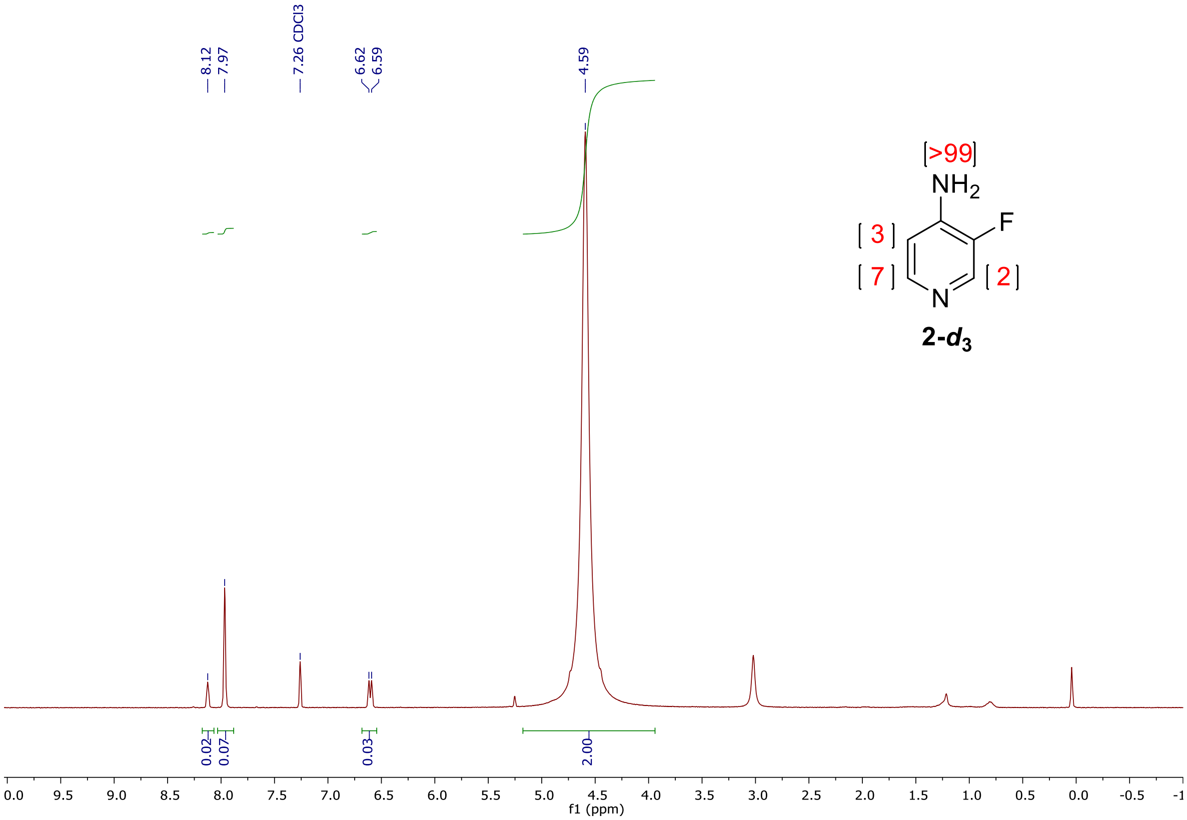

{[1]H NMR} spectrum of 3-fluoro-4-aminopyridine-2,5,6-*d*3 (**2-*d*3**).

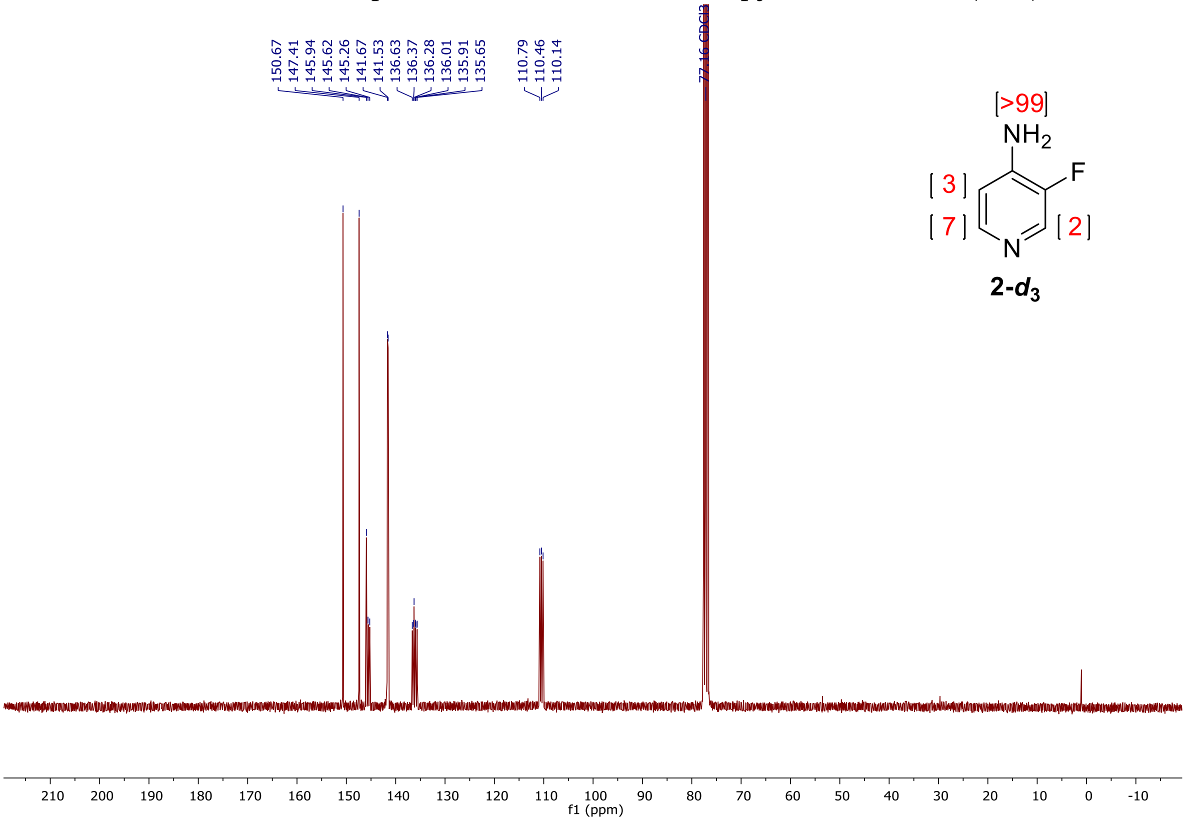

{[13]C NMR} spectrum of 3-fluoro-4-aminopyridine-2,5,6-*d*3 (**2-*d*3**).

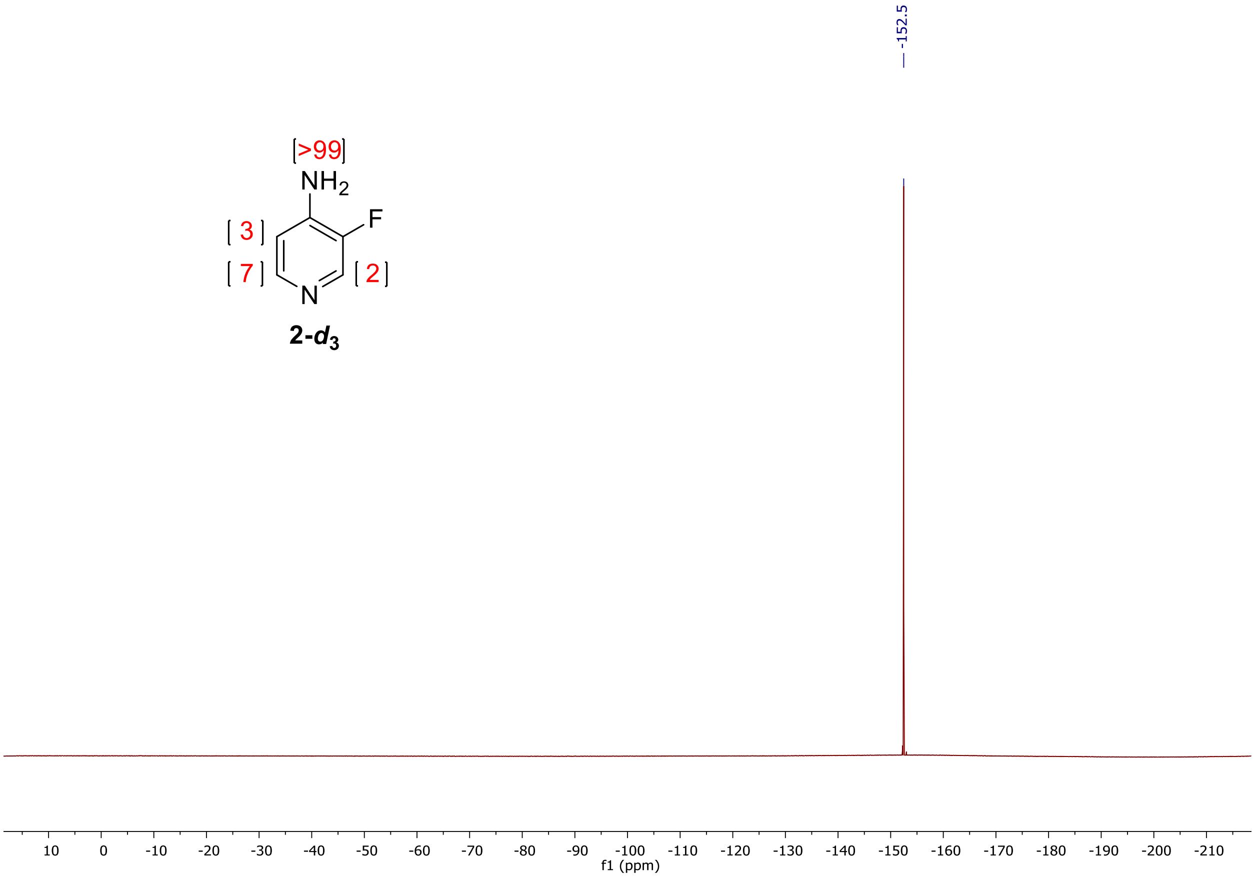


{$^{19}$F NMR} spectrum of 3-fluoro-4-aminopyridine-2,5,6-*d*$_3$ (**2-*d*$_3$**).

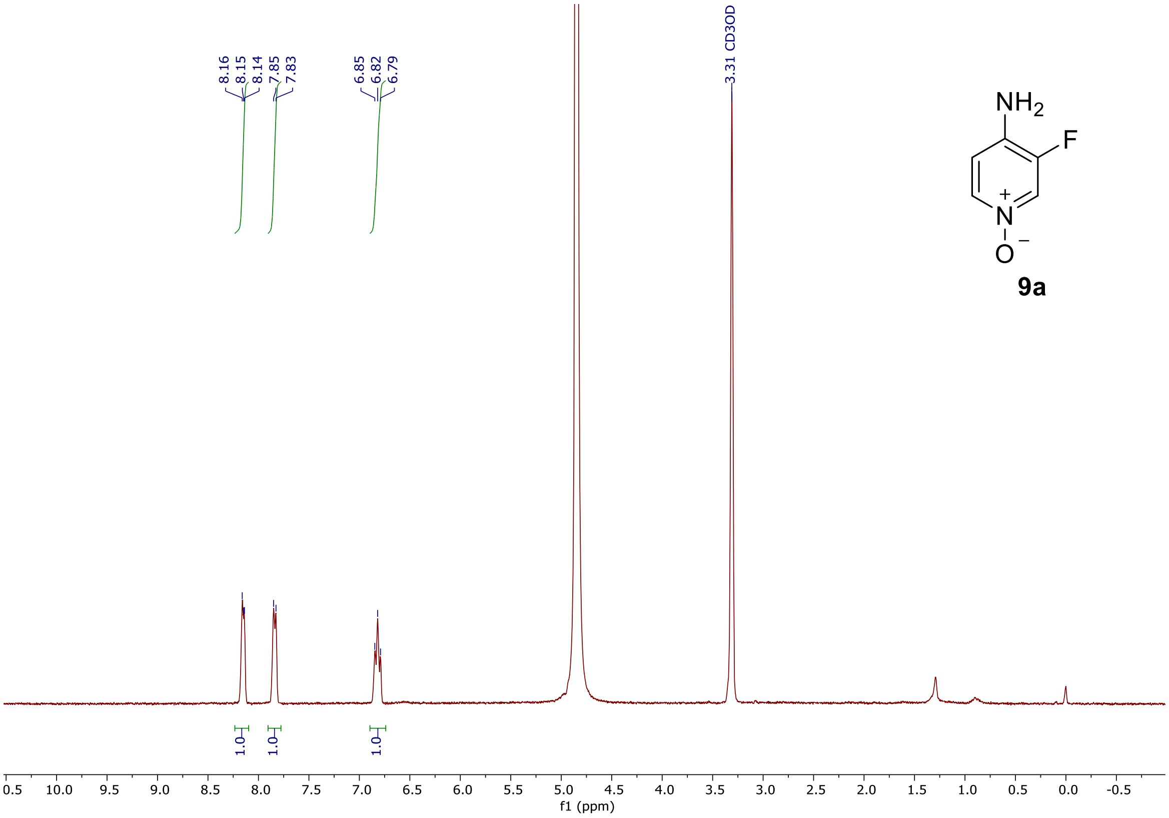


{[1]H NMR} spectrum of 3-fluoro-4-aminopyridine 1-oxide (**9a**).

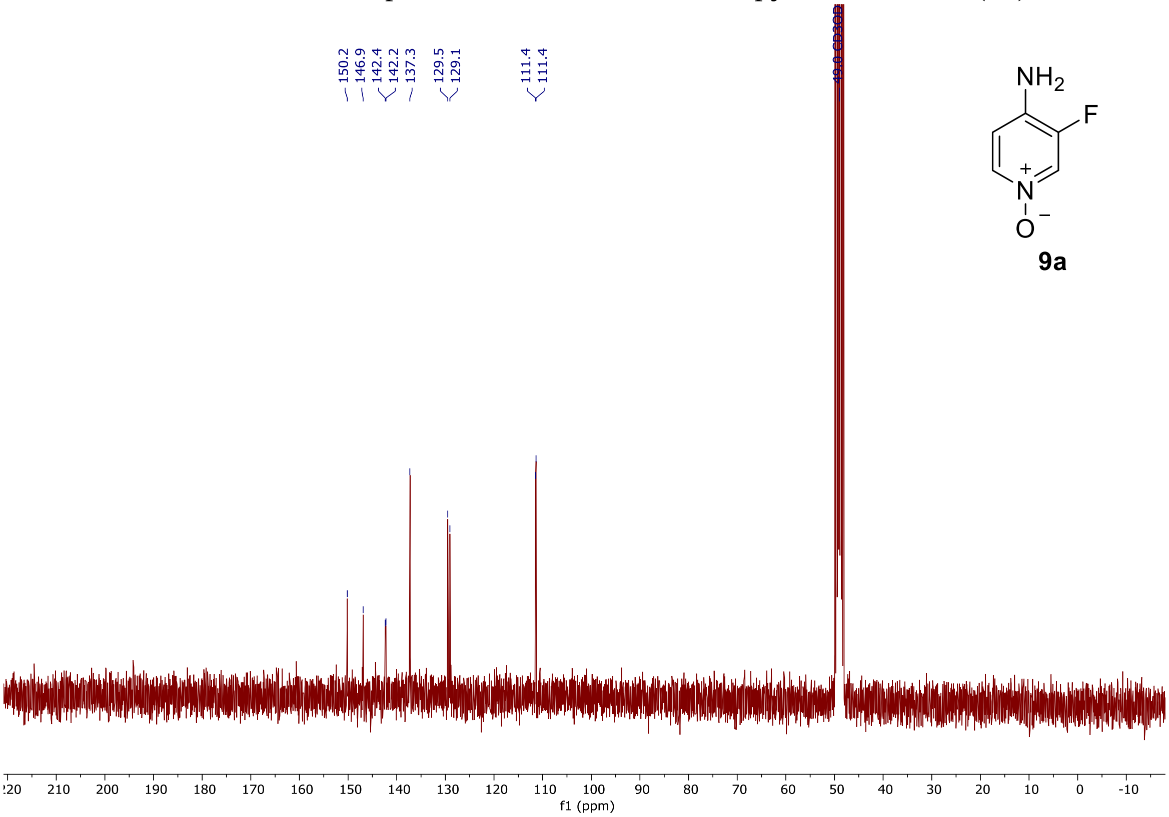


{[13]C NMR} spectrum of 3-fluoro-4-aminopyridine 1-oxide (**9a**).

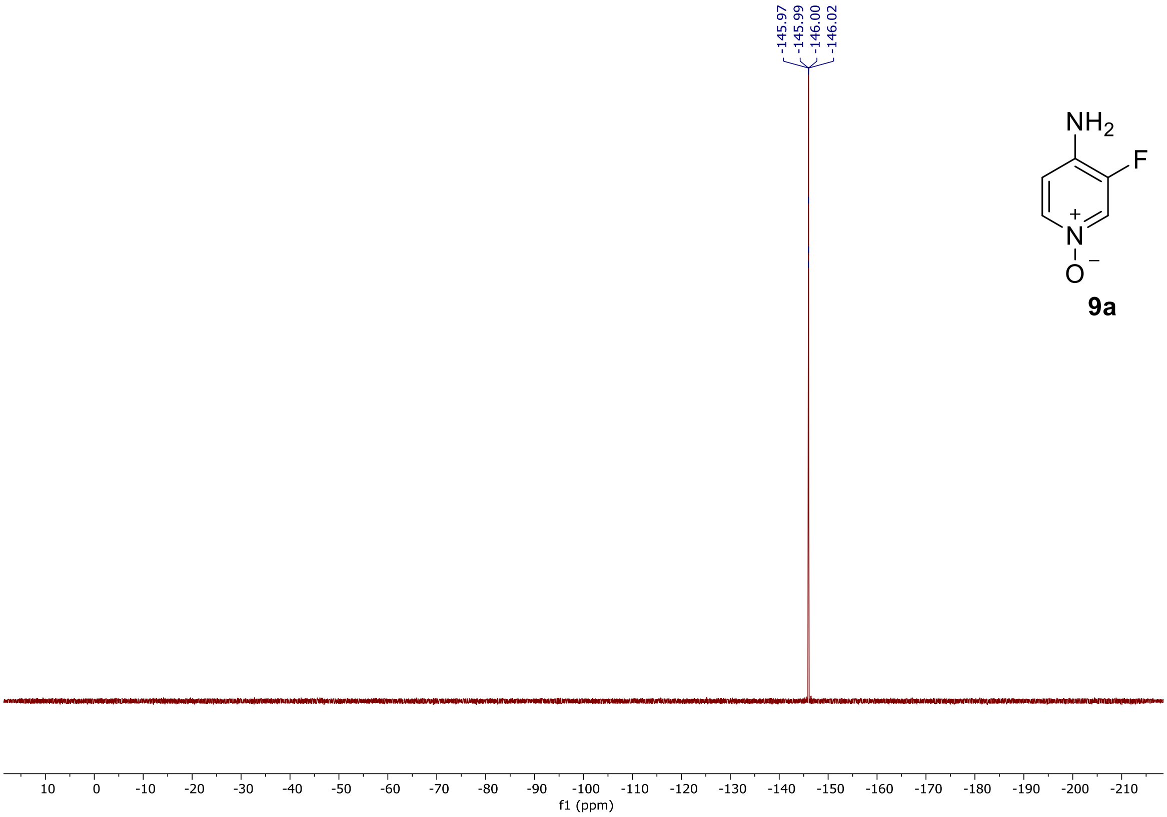


{$^{19}$F NMR} spectrum of 3-fluoro-4-aminopyridine 1-oxide (**9a**).
